# A mitochondrial genome phylogeny of voles and lemmings (Rodentia: Arvicolinae): evolutionary and taxonomic implications

**DOI:** 10.1101/2021.02.23.432437

**Authors:** Natalia I. Abramson, Semyon Yu. Bodrov, Olga V. Bondareva, Evgeny A. Genelt-Yanovskiy, Tatyana V. Petrova

## Abstract

Arvicolinae is one of the most impressive placental radiations with over 150 extant and numerous extinct species that emerged since the Miocene in the Northern Hemisphere. The phylogeny of Arvicolinae has been studied intensively for several decades using morphological and genetic methods. Here, we sequenced 30 new mitochondrial genomes to better understand the evolutionary relationships among the major tribes and genera within the subfamily. The phylogenetic and molecular dating analyses based on 11,391 bp concatenated alignment of protein-coding mitochondrial genes confirmed the monophyly of the subfamily. While Bayesian analysis provided a high resolution across the entire tree, Maximum Likelihood tree reconstruction showed weak support for the ordering of divergence and interrelationships of tribal level taxa within the most ancient radiation. Both the interrelationships among tribes Lagurini, Ellobiusini and Arvicolini, comprising the largest radiation and the position of the genus *Dinaromys* within it also remained unresolved. For the first time complex relationships between genus level taxa within the species-rich tribe Arvicolini received full resolution. Particularly *Lemmiscus* was robustly placed as sister to the snow voles *Chionomys* in the tribe Arvicolini in contrast with a long-held belief of its affinity with Lagurini. Molecular dating of the origin of Arvicolinae and early divergences obtained from the mitogenome data were consistent with fossil records. The mtDNA estimates for putative ancestors of the most genera within Arvicolini appeared to be much older than it was previously proposed in paleontological studies.

## Introduction

Reconstructing the phylogeny of a taxonomic group that emerged during the rapid species diversification is the major challenge in evolutionary biology. When a small number of molecular markers are considered in the study, unresolved bush-like trees or polytomies are obtained. The subfamily Arvicolinae Gray, 1821 (Rodentia: Cricetidae) consisting of voles, lemmings and muskrats, provide a good example for the research of systematics and taxonomy of fast species radiations within mammals. Arvicolinae is a highly diverse, youngest and fast-evolving group within the order Rodentia. During the rapid explosive radiation and diversification that started in the Late Miocene according to the fossil records, voles and lemmings occupied all types of landscapes within the temperate and cold climate biomes of the Northern Hemisphere. The morphological evolutionary history of the group is perfectly documented using the rich fossil records upon a number of extinct species described. Modern global fauna of Arvicolinae contains 151-162 recent species grouped into 28 genera [1, 2], and new species are constantly being discovered and described. Similarly to other taxa that experienced rapid adaptive radiation, phylogenetic reconstruction of this group faces several principal methodological problems, as ecological convergence and homoplasy severely limit the use of morphological traits in phylogenetic analysis.

Phylogeny of Arvicolinae has been explored using both morphological and genetic methods, allowing comparisons of reconstruction from different datasets for further cross-validation. Application of molecular phylogenetic methods resulted in a series of revisions of phylogenetic relationships and taxonomic structure of several genera and species [3–12] and references therein. Reconstruction of Arvicolinae phylogeny using nuclear genes *GHR* and *LCAT* demonstrated three successive waves of adaptive radiations in the evolutionary history of the group [7]. The first radiation wave most plausibly took place in Late Miocene and marked the emergence of muskrats (Ondatrini), lemmings (Lemmini and Dicrostonychini) and long-clawed mole voles (Prometheomyini). Second radiation wave is characterized by a divergence of the ancestors of modern Clethrionomyini. The third radiation wave included formation of the steppe lemmings (Lagurini), mole voles (Ellobiusini) and the richest species group - Arvicolini [7]. The branching order both within the first and the last radiation waves also remained unresolved since all attempts to untangle complex phylogenetic relationships within subfamily were made with the use of only a few mitochondrial or nuclear markers, or the study included an insufficient number of taxa in the analysis [3,5,7,9,10,12–18].

With the upcoming epoch of genomic studies, it is obvious that further breakthrough in the study of the extremely complex evolutionary history of the subfamily Arvicolinae can only be achieved by switching from the “gene-centric” approach to the analysis of genomic datasets, yet comprehensive sampling of taxa is also important. Analyses of complete mitochondrial genomes have been successfully used to reconstruct robust phylogenies in many animal groups [19–23] and other. The number of published mitogenomes of voles, lemmings and muskrats is permanently increasing [24–38] and other, and data on nearly one-fifth of the total species diversity (ca. 30 species) is already available.

Though molecular studies during the last decades have considerably extended and refined our knowledge of the pattern and timescale of arvicoline phylogeny, there are important issues that remain to be elucidated. In this study, we were aimed on estimating the phylogeny of Arvicolinae using complete mitogenomes generated using high-throughput sequencing. By significantly increasing the number of newly sequenced mitogenomes representing major tribes of voles and lemmings, we implement the phylogenetic and molecular dating analysis on a dataset consisting of almost all living genera within the subfamily. The following questions were specifically addressed during the study: (1) the order of divergence and interrelationships of taxa within the first, most ancient radiation, (2) the interrelationships of three tribe level taxa Lagurini, Ellobiusini and Arvicolini (3) relative phylogenetic placement of genera *Dinaromys* Kretzoi, 1955 and *Lemmiscus Thomas, 1912*, (4) tangled interrelationships of genera and subgenera in the most speciose tribe Arvicolini, (5) the position of *Agricola agrestis* Linnaeus, 1761 and *Iberomys cabrerae* Thomas, 1906, and (6) the timing of arvicoline divergences.

## Material and methods

### Taxonomic sampling

Fifty-eight species of Arvicolinae, belonging to 27 genera and all the tribe level taxa, as well as six outgroup taxa were used in this study. Complete mitochondrial genomes for 30 Arvicolinae species were sequenced in the current study (including 15 species belonging to the Arvicolini tribe, one for Dicrostonychini, two for Lagurini, three for Lemmini, six species belonging to Clethrionomyini and three crucial species without stable taxonomic position: *Prometheomys schaposchnikowi* Satunin, 1901 (Prometheomyini), *Dinaromys bogdanovi* Martino, 1922 and *Lemmiscus curtatus* Cope, 1868). For 28 species belonging to Arvicolini, Ellobiusini, Clethrionomyini, Dicrostonychini and Ondatrini tribes sequences were available in the NCBI database. The detailed information, GenBank accession numbers, and the voucher IDs for new sequences are given in S1 Table. Hereinafter, we use the taxonomic classification following Gromov & Polyakov [39], Musser & Carleton [1], Abramson & Lissovsky [40] with amendments made in result of the current study.

### DNA isolation, NGS library preparation and sequencing

Muscle tissue samples of fresh specimens were collected between 1996-2019 years and stored in 96% ethanol at -20 degrees Celcius in a tissue and DNA collection of the Group of molecular systematics of mammals (Zoological Institute RAS). Historic specimen of *Lemmiscus curtatus* (sampled in 1927) was obtained from the collection of the laboratory of theriology (Zoological Institute RAS), see S1 Table for details.

Homogenization of tissues was performed using the Qiagen TissueLyser LT (Qiagen). For the most samples, genomic DNA was extracted using the Diatom DNA Prep 200 (Isogen, Russia) except for the *L. curtatus* museum specimen. To reduce the potential contamination, all manipulations with the *L. curtatus* were carried out in a separate laboratory room isolated from post-PCR facilities, predominantly being used for studies of historic samples from the collection of Zoological Institute. All the working surfaces, instruments and plastics were sterilized with UV light and chloramine-T. DNA from the museum skin sample (2 × 2 mm piece from the inner side of the lip, dissected by a sterilized surgical blade) was isolated using the phenol-chloroform extraction method according to a standard protocol [41]. PCR was prepared using a PCR workstation (LAMSYSTEMS CC, Miass, Russia).

The following ultrasound fragmentation of the total genomic DNA was implemented using Covaris S220 focused ultrasonicator instrument (Covaris). The resulting fragmented DNA was purified and concentrated using paramagnetic bead-based chemistry AMPure XP (Beckman-Coulter) using standard workflow. DNA concentration was evaluated using a Qubit fluorometer (Thermo Fisher Scientific).

NGS libraries were prepared using the NEBNext Ultra II DNA Library Prep Kit for Illumina (New England Biolabs). The resulting PCR products were purified and concentrated using AMPure XP beads (Beckman-Coulter). The concentration of samples was measured using a Qubit fluorometer, and quality control of the libraries was implemented using Bioanalyzer 2100 instrument and the DNA High Sensitivity kit (Agilent). Sequencing was performed on an Illumina HiSeq 4000 system, resulting in pair-end reads of 75bp. DNA quality was checked with Qubit, the final distribution of lengths of the libraries adapter content checking was conducted using Bioanalyzer2100 (Agilent). DNA extraction (except the museum specimen of *L. curtatus*), library preparation and sequencing were performed using resources of the Skoltech Genomics Core Facility (https://www.skoltech.ru/research/en/shared-resources/gcf-2/).

### Read processing, mitogenome assembly and annotation

The quality of raw reads was evaluated using FastQC [42], and parts with the quality score below 20 were trimmed using Trimmomatic-0.32 [43]. Bowtie 2.3.5.1 [44] was used to filter reads with contamination. Complete mitochondrial genomes of other Arvicolinae were used as reference sequences. Also, this was made for the museum specimen to enrich reads with mitochondrial DNA.

Nucleotide misincorporation patterns that can often be observed during the studies of ancient or old museum sample DNA as a result of post-mortem DNA damage in reads from *L. curtatus* were achieved using mapDamage 2.0 [45].

Complete mitochondrial genome was assembled using *plasmidSPAdes* [46] with default settings. The resulting contigs were filtered by length, the most similar in size to mitochondrial DNA were selected (size about 16 kb for mammals). The contigs were annotated using the online-server MITOS [47] http://mitos2.bioinf.uni-leipzig.de/index.py, with default settings and the vertebrate genetic code for mitochondria.

Gene boundaries were checked and refined by alignment against 28 published mitogenome sequences of Arvicolinae (see details in S1 Table). All positions of low quality, low coverage, as well as fragments that greatly differed from the reference Arvicolinae mitochondrial genomes, were replaced by N manually. Assembled sequences of protein coding genes (PCGs) were checked for internal stops manually. All assembled and annotated mitogenomes have been deposited in GenBank (S1 Table).

### Sequence alignment

The 30 newly obtained mitochondrial genomes were compared with 28 earlier published Arvicolinae mitogenomes mined from NCBI (see accession numbers in S1 Table), including and five mitogenomes obtained by us earlier [48–50]. All mitochondrial genomes were aligned with Mauve (http://darlinglab.org/mauve/mauve.html) in Geneious Prime 2019.1 (Biomatters Ltd.).

In several studies, it has been convincingly shown that protein-coding sequences may have a strong resolving power for inferring phylogenetic interrelationships and divergence time estimates derived from PCG may be quite accurate [22,51,52]. We used this approach, however complete mt genomes will serve as the starting point for further analyses. For the subsequent analyses, the concatenated alignment of 13 PCGs using MAFFT version 7.222 [53] was produced.

Third codon position has previously been shown to bias phylogenetic reconstructions [54]. The phylogeny on a smaller dataset of Arvicolinae turned out to be very poorly resolved with the exception of third codon position [50]. So we masked transitions in 3rd codon position by RY-coding (R for purines and Y for pyrimidines) as described in Abramson et al. [50].

Thus, two datasets were subsequently analyzed — total alignment of 13 PCGs, where all three codon positions were considered (with a length of 11,391 bp) and RY-coded alignment with transitions in third codon position masked.

### Analysis of base composition

The base composition was calculated in Geneious Prime 2019.1 (Biomatters Ltd.). The strand bias in nucleotide composition was studied by calculating the relative frequencies of C and G nucleotides (CG3 skew = [C − G]/[C + G]) [22,55,56]. Both analyses were calculated using full-length mitogenomes.

The PCG-alignment of 64 mitochondrial genomes was used to calculate relative frequencies of four bases (A, C, G and T) at each of three codon positions in MEGA X [57]. The 12 variables, each representing base frequency in first, second or third position, were then summarized by a Principal Component Analysis (PCA) using the PAST v.4.04 [58].

### Saturation tests

The presence of phylogenetic signal was assessed with a substitution saturation analysis using the Xia test [59] in the DAMBE 7.2.1 software [60] for the whole alignment of the PCG dataset and 13 separate genes following the procedure described by Xia & Lemey [61], particularly when (a) 1st and 2nd codon position considered and 3rd position is masked from the alignment, and (b) when only 3rd codon position is included in the analysis. The analysis is based on Index of substitution saturation - *Iss*, and *Iss.c* is the critical value at which the sequences begin to fail to recover the true tree).

Once *Iss.c* is known for a set of sequences, then we can calculate the Iss value from the sequences and compare it against the *Iss.c*. If Iss value exceeds the *Iss.c*, we can conclude that the sequence dataset consists of substitution saturation and cannot be used for further phylogenetic reconstruction.

The proportion of invariant sites was specified for tests considered 1st and 2nd codon positions. The analysis was performed on a complete alignment with all sites considered. Additional analysis of saturation for each of the PCG was estimated using R packages *seqinr* [62] and *ape* [63]. P-distances were plotted against K81 distances for transitions and transversions of each codon position.

### Phylogenetic analyses

We used PartitionFinder 2.1.1 [64] applying AICc and “greedy” algorithm, when an analysis is based on the a priori features of the alignment, to select the optimal partitioning scheme for each dataset. Our analysis started with the partitioning by codon positions within PCG fragments, each treated as a unique partition. For the complete 13 PCG alignment, GTR+I+G model was suggested almost for all the partitions except *ND6* 3rd codon position, for which the TRN+I+G model was selected. For the alignment with RY-coded 3rd codon position, two partitions were suggested - 1+2nd and 3rd codon positions with GTR+I+G and GTR+G models respectively.

Phylogenetic reconstructions using Maximum Likelihood (ML) and Bayesian Inference (BI) analyses were performed on both complete and RY-masked datasets partitioned as suggested with PartitionFinder. Trees were rooted by six sequences of Cricetinae: *Akodon montensis* Thomas, 1913, *Peromyscus megalops* Merriam, 1898, and four species of hamsters from genus *Cricetulus* Milne-Edwards, 1867 (S1 Table).

Maximum Likelihood (ML) analysis was performed using IQ-TREE web server [65] with 10,000 ultrafast bootstrap replicates [66]. Bayesian Inference (BI) analysis was performed in MrBayes 3.2.6 [67]. Each analysis started with random trees and performed two independent runs with four independent Markov Chain Monte Carlo (MCMC) for 10 million generations with sampling every 1,000th generation, the standard deviations of split frequencies were below 0.01; potential scale reduction factors were equal to 1.0; stationarity was examined in Tracer v1.7 [68]. A consensus tree was constructed based on the trees sampled after the 25% burn-in.

We also conducted ML analysis for each PCG separately (with partitions by codon positions and models supposed with IQ-TREE). *Hyperacrius fertilis* True, 1894 sequence was excluded from the *ND4* alignment since this gene was highly fragmented [50]. The mitogenome of *Craseomys rufocanus* Sundevall, 1846 (accessed from GenBank) completely lacked the *ND6* sequence (S2 Table), so this species was excluded from the analysis for this gene.

### Divergence dating

Divergence times were estimated on the CIPRES Science Gateway [69] with Bayesian approach implemented in BEAST v.2.6.2 [70] using both complete PCG dataset and the one in which the transitions in the third codon position were masked with RY-coding. Datasets were partitioned according to the recommendations of PartitionFinder. All site model parameters were chosen for separate partitions with corrected Akaike’s information criterion (AICc) in JMODELTEST 2.1.1 [71]. Eight fossil calibrations were used (S3 Table). Lognormal prior distributions were applied to all the calibrations with offset values and 95% HPD intervals based on first appearance data (FAD) and stratigraphic sampling downloaded from the Paleobiology Database on 01.12.2020 using the parameters “Taxon = fossil species, Timescale = FAD” (S3 Table).

BEAST analyses under the birth-death process used a relaxed lognormal clock model and the program’s default prior distributions of model parameters. Each analysis was run for 100 million generations and sampled every 10 000 generations. The convergence of two independent runs was examined using Tracer v1.7 [68], and combined using LogCombiner, discarding the first 25% as burn-in. Trees were then summarized with TreeAnnotator using the maximum clade credibility tree option and fixing node heights as mean heights. Divergence time bars were obtained automatically in FigTree v1.4.3 (http://tree.bio.ed.ac.uk/software/figtree/) from the output using the 95% highest posterior density (HPD) of the ages for each node.

## Results

### Mitochondrial genome assembly and annotation

We sequenced, assembled and annotated mitochondrial genomes for the 30 new taxa of Arvicolinae. The mapDamage analysis implemented on the raw reads of *Lemmiscus curtatus* (S1 Fig) showed a low variation of deamination misincorporations values. C to T misincorporations varied from 10 to 15%, G to A from 10 to 12% and was equal to the results of similar studies [72]. Since the relative level of observed misincorporations was not significantly different from the other substitution variants, the mitogenome of *L. curtatus* was assembled using the same pipeline as for the rest of taxa.

The assembled mitogenomes, circular double-stranded DNA of the same organization as in other mammals, contained 13 PCGs, 22 transfer RNAs (tRNA), two ribosomal RNAs (rRNAs), and a non-coding region corresponding to the control region (D-loop). Nine genes (*ND6* and eight tRNAs) were oriented in the reverse direction, whereas the others were transcribed in the forward direction. All the assembled mitogenomes contained all the genes listed above, but in some species demonstrated incomplete gene sequences (see S2 Table for details).

Mitochondrial genome sequences were deposited in GenBank under accession numbers indicated in S1 Table. In the subsequent analyses, the PCG dataset, containing 11,391 bp was used.

### Variation in base composition

Comparison of base composition calculated using the alignment of full-length mitogenome sequences of Arvicolinae showed that mitochondrial genomes of taxa from tribes Clethrionomyini (28.36% С) and Ellobiusini (28.7% C) have the highest GC-content. Arvicolini, Dicrostonychini and Lemmini had slightly smaller values: 27.69, 28.03 and 27.77% C respectively. Lagurini were found to have the most AT-skewed base composition of mitogenomes: 31.20 and 31.30% of adenine, respectively (S1 Table). Lagurini and Arvicolini also demonstrated the highest GC-skew values (-0.32 in both cases). Ellobiusini and Clethrionomyini occupied an intermediate position in terms of it with -0.33 and -0.34, respectively. Dicrostonychini and Lemmini with equal value -0.35 have the smallest GC-skew values.

The base composition (frequency of the nucleotides A, C, G, and T) was further analyzed at the three codon positions in the concatenated alignment of PCGs for each species separately (S1 Table). The 12 variables measured for 64 taxa were summarized by a PCA, based on the first two components, which contributed 73.7% and 18.8% of the total variance, respectively (Fig 1). Most of the observed variation was related to the percentage of base composition in a third codon position. The first component demonstrated a high positive correlation (0.98) with the percentage of C3 (percentage of cytosine in the third position) and a high negative correlation (-0.93) with T3 (thymine in the third position). The second component positively correlated with G3 (0.67) and negatively correlated with A3 (-0.88). Most of the Arvicolinae formed a compact group on the PCA graph. Among the highly dissimilar to the main group were almost all *Cricetulus* species. The two rest outgroup taxa, *C. kamensis Satunin, 1903* and *Akodon montensis* grouped with Arvicolinae, and *A. montensis* showed similar base composition to *Hyperacrius fertilis*. Among Arvicolinae, the most dissimilar base composition was observed in *Mynomes longicaudus* Merriam, 1888, *Chionomys gud* Satunin, 1909 and *Arvicola amphibius* Linnaeus, 1758, showing higher than group average percentage of T3 and lower than group average percentage of C3. The mitochondrial genome of *Ondatra zibethicus* Linnaeus, 1766 was characterized by the highest percentage of adenine in the third position (45.7%) compared to other Arvicolinae. *Ellobius lutescens* Thomas, 1897 demonstrated the highest percentage of cytosine in third position (37.1%) among the complete PCG dataset (Fig 1).

**Fig 1.**
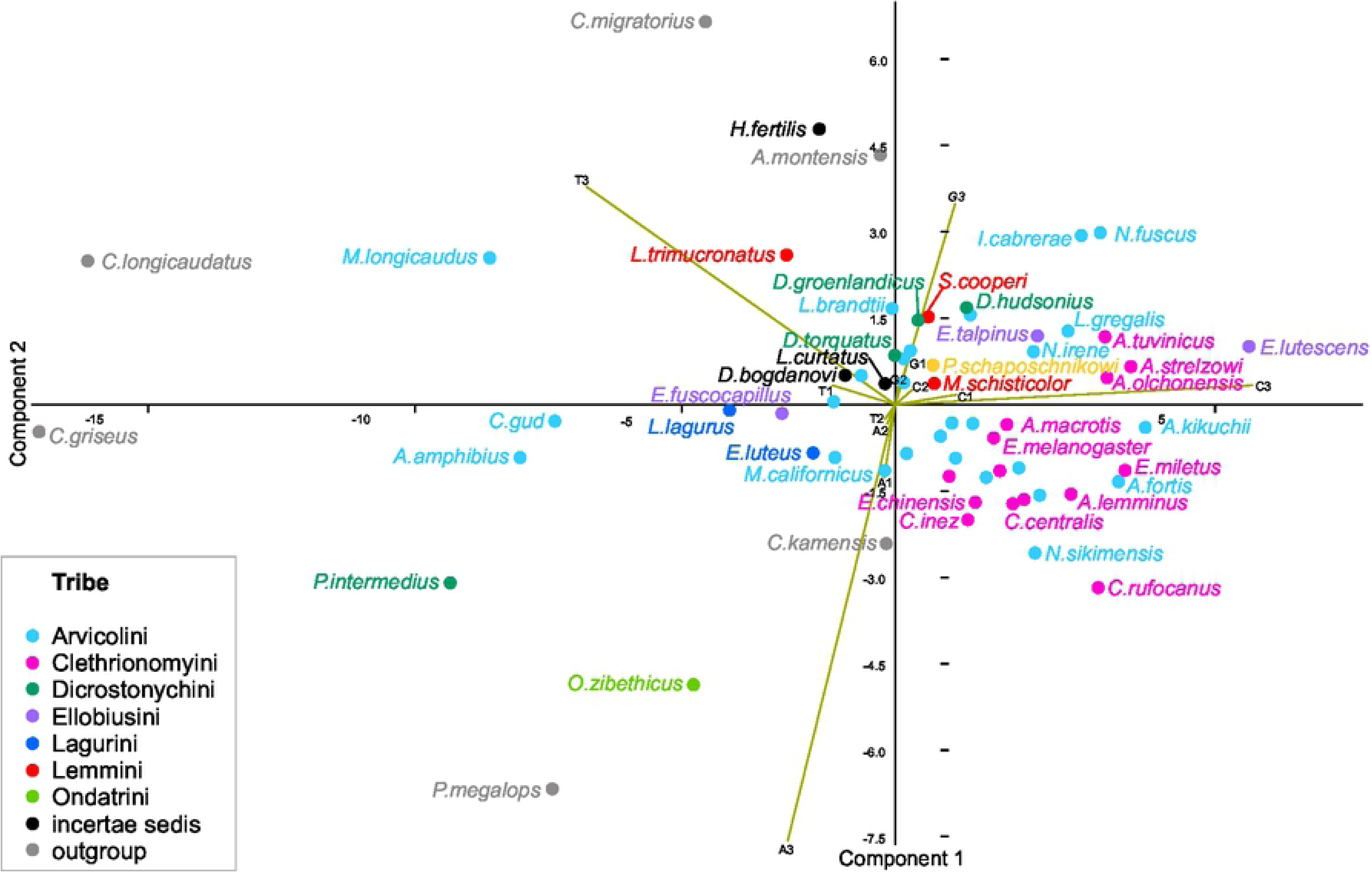
Base composition in mitochondrial PCG of Arvicolinae. The frequency of the four bases (A, C, G, and T) at each codon position (first, second and third) in concatenated alignment was used as 12 variables for PCA. Tribes are indicated by colours.

### Substitution saturation analysis

Substitution saturation decreases phylogenetic information contained in the sequences and plagues the phylogenetic analysis involving deep branches.

According to the analysis implemented in DAMBE software (S4 Table), the observed *Iss* saturation index was significantly (P<0.0001) lower than critical *Iss.c* value for both symmetrical and asymmetrical topology tests indicating the lack of saturation in the studied Arvicolinae dataset.

The results of saturation plots for separate genes show the same pattern of negligible saturation. As a result for all 13 PCGs, no significant saturation for the 1st and 2nd codon position, and they are all suitable for the phylogenetic inference. *CYTB*, *ND1* and *ND6* show the same for 3rd codon position in contrast with *ND2*, *ND3*, *ND4*, *ND4L.* For other genes there was no significant saturation even for the 3rd codon position considering symmetrical topology for more than 32 OTUs (number of operational taxonomic units, S4 Table).

### Time-calibrated mitochondrial genome phylogeny of Arvicolinae

The maximum-likelihood (ML) and Bayesian inference (BI) trees reconstructed using complete and RY-coded alignments of PCGs had similar topology (Fig 2). Overall, ML analysis demonstrated lower node supports compared to BI analysis. In total 70% of the nodes were highly supported by ML and BI, with Bayesian probabilities BP>0.95 and ML bootstrap support BS>95 (Fig 2, nodes with a black dot).

**Fig 2.**
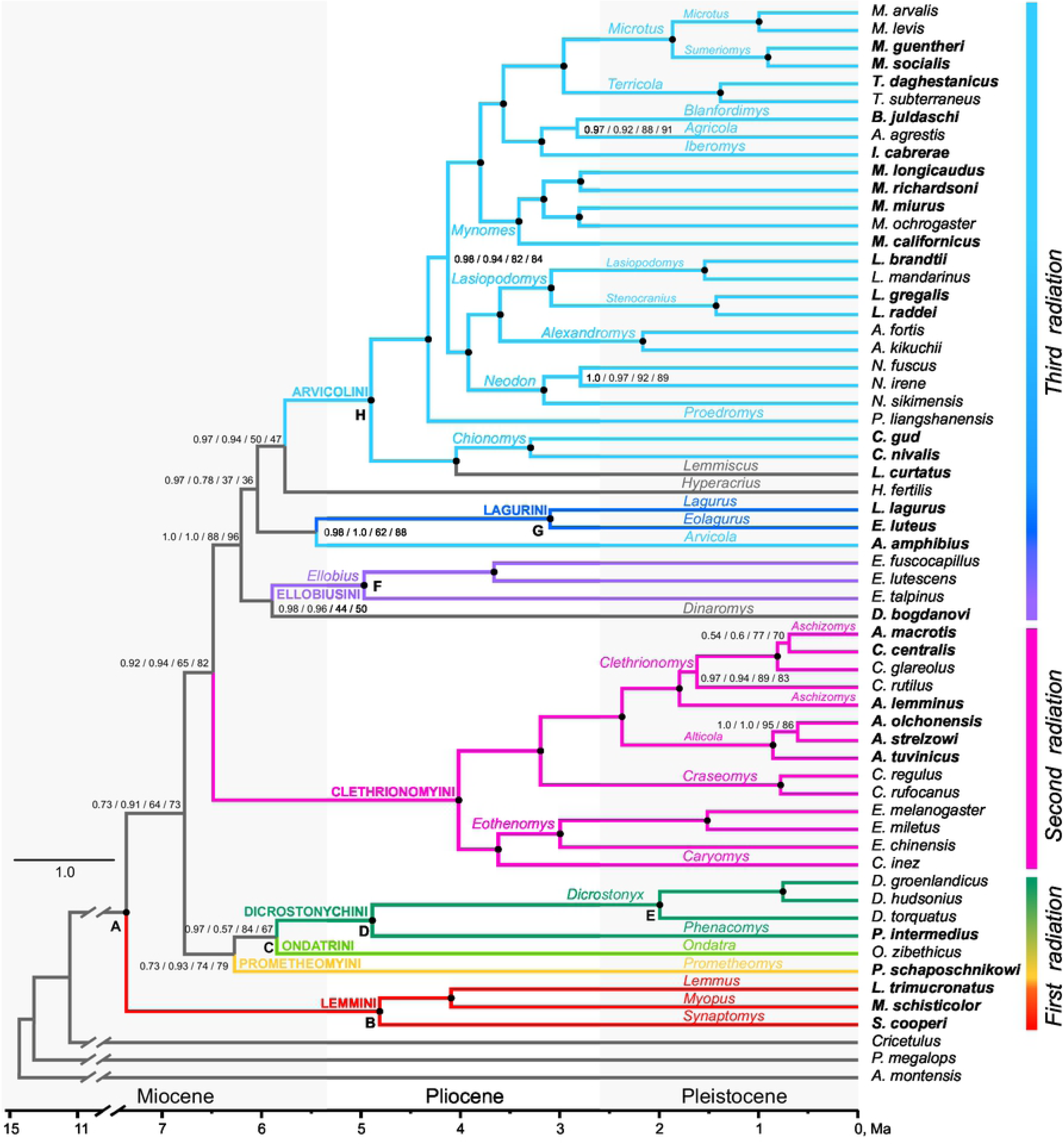
Time-calibrated mitochondrial phylogeny of Arvicolinae. Node labels display the following supports: BI complete / BI RY-coded 3rd codon position / ML complete / ML RY-coded 3rd codon position. Black circles show nodes with 0.95-1.0 BI and 95-100 ML support. All letters at nodes correspond to fossil constraints in S2 Table. Traditional tribal designations are also given above the branches and corresponding branches distinguished by different colors.

The monophyly of subfamily Arvicolinae was strongly supported by BI and ML analyses. However, several nodes, predominantly the internal nodes representing deeper phylogenies, which were highly supported by Bayesian analysis, did not receive high BS values. The divergence time between Arvicolinae and Cricetinae was estimated as Late Miocene, ca. 11.31 / 10.7 Ma, based on the complete and RY-coded alignments respectively (Table 1).

**Table 1.** Divergence time estimates for the major lineages within the subfamily Arvicolinae.

| TMRCAs | Complete | RY-masked |
| --- | --- | --- |
| Arvicolinae + Cricetinae | **11.31** (9.48-13.3) | **10.7** (8.42-13.31) |
| Arvicolinae | **7.36** (7.04-7.78) | **7.33** (7.05-7.73) |
| Lemmini | **4.81** (3.68-5.97) | **4.37** (3.31-5.71) |
| Dicrostonychini + Ondatrini | **5.85** (5.15-6.56) | **5.54** (4.41-6.59) |
| Dicrostonychini | **4.89** (4.08-5.7) | **4.49** (3.23-5.86) |
| Clethrionomyini | **4.02** (3.33-4.72) | **4.46** (3.35-5.64) |
| (Ellobiusini + *Dinaromys*) / (Arvicolini + Lagurini) | **6.2** (5.65-6.76) | **6.11** (5.17-6.92) |
| Ellobiusini | **4.97** (4.21-5.69) | **4.58** (3.42-5.68) |
| Lagurini | **3.1** (2.59-3.75) | **3.05** (2.56-3.75) |
| *Hyperacrius* / Arvicolini | **5.76** (5.2-6.34) | **5.79** (4.88-6.63) |
| Arvicolini s.str.* | **4.9** (4.33-5.47) | **5.02** (4.12-5.89) |
| *Chionomys* + *Lemmiscus* | **4.04** (3.3-4.74) | **4.28** (3.11-5.4) |
| *Chionomys* | **3.29** (2.5-4.04) | **3.45** (2.13-4.67) |
| *Proedromys* / | **4.32** (3.81-4.86) | **4.52** (3.69-5.33) |
| *Microtus* ** (*Microtus*, *Sumeriomys*, *Terricola*,<br>*Blanfordimys*, *Agricola*, *Iberomys*) | **3.8** (3.31-4.3) | **3.87** (3.07-4.63) |
| *Mynomes* | **3.41** (2.89-3.91) | **3.32** (2.48-4.11) |
| *Microtus* + *Terricola* | **2.96** (2.46-3.5) | **3.18** (2.35-3.95) |
| *Microtus* | **1.87** (1.43-2.33) | **2.06** (1.31-2.8) |
| *Terricola* | **1.38** (0.89-1.93) | **1.46** (0.7-2.32) |
| *Iberomys* + (*Agricola* + *Blanfordimys*) | **3.18** (2.66-3.71) | **3.11** (2.23-3.99) |
| *Agricola* + *Blanfordimys* | **2.82** (2.26-3.35) | **2.64** (1.69-3.54) |
| *Neodon* + (*Alexandromys* + *Lasiopodomys*) | **3.92** (3.4-4.4) | **4.12** (3.31-4.9) |
| *Neodon* | **3.16** (2.58-3.76) | **3.3** (2.43-4.23) |
| *Alexandromys* + *Lasiopodomys* | **3.6** (3.12-4.09) | **3.65** (2.86-4.44) |
| *Lasiopodomys* | **3.09** (2.59-3.56) | **3.07** (2.27-3.85) |
| *Alexandromys* | **2.16** (1.55-2.81) | **2.2** (1.27-3.22) |

Mean node ages marked in bold and 95% highest posterior density intervals (in brackets) in million years ago (Ma) estimated with two PCG datasets - complete and with RY-masked 3rd codon position.
\* excluding *Arvicola* and *Hyperacrius*
\*\* sensu Pardinas et al., 2017.

The earliest radiation of the proper arvicolines (tribes Lemmini, Prometheomyini, Ondatrini and Dicrostonychini) dates back to the Late Miocene with mean at 7.36 / 7.33 Ma. Despite the high node support for the nodes marking tribes Lemmini and Dicrostonychini, the basal part of the phylogenetic tree remains unresolved and represents a polytomy with several nodes not receiving significant BI and ML support. Analysis based on the PCG dataset where the third codon position was not masked with RY-coding, indicated significantly high Bayesian support for node C, combining Ondatrini and Dicrostonychini (Fig 2). The time to MRCA of node C is about 5.85 / 5.54 Ma and the MRCA of proper Dicrostonychini at 4.89 / 4.49 Ma.

The tribe Clethrionomyini representing second radiation of Arvicolinae received high BI and ML support, and nodes within the clade were also highly supported. The MRCA for Clethrionomyini dates back to 4.02 / 4.46 Ma.

The cluster containing tribes Ellobiusini, Lagurini, Arvicolini and genera *Dinaromys, Arvicola* Lacepede, 1799 and *Hyperacrius* Miller, 1896, i.e. the third radiation of Arvicolinae, was robustly supported by BI using both alignments and received reliable support by ML only with RY-masked alignment (Fig 2). Within this cluster, nodes marking tribes were highly supported by BI and ML. At the level of terminal branches within this cluster *Dinaromys bogdanovi, Hyperacrius fertilis and Arvicola amphibius* were the only to lack a certain phylogenetic position. *D. bogdanovi* grouped with Ellobiusini showing high BI support and no ML support. The water vole, *Arvicola amphibius, clustered* with Lagurini (high BP and no BS support) thus being paraphyletic to Arvicolini. The sagebrush vole *Lemmiscus curtatus* was sister to snow voles, *Chionomys gud* and *C. nivalis* Martins, 1842 with a robust support obtained in all analyses. The cluster of *Chionomys* Miller, 1908 and *Lemmiscus* was the earliest derivative in the highly supported group uniting all known vole genera of Arvicolini tribe. Arvicolini *sensu stricto* (excluding *Arvicola*) was fully resolved: both node H marking the whole tribe and all nodes within the tribe received robust support in ML and BI analyses.

The estimated time of the largest radiation event within the subfamily and TMRCA for the trichotomy Arvicolini - Ellobiusini - Lagurini dates back to 6.2 / 6.11 Ma. All following divergence events within this radiation according to the obtained estimates took place very close to each other, the 95% HPD of the diverging branches leading to MRCA of existing tribes are highly overlapping. Thus the date estimate to the MRCA of Ellobiusini was 4.97 / 4.58 Ma. The MRCA of the Lagurini tribe is around 3 Ma (Fig 2, Table 1). The estimate for the earliest split within the Arvicolini tribe radiation (not including *Arvicola* and *Hyperacrius*) with all recent genera is about 4.9 / 5.02 Ma, that coincides with the onset of Pliocene period, whereas the major part of recent genera, excluding early derivating *Chionomys*, *Lemmiscus* and *Proedromys* Thomas, 1911, according to obtained estimates appear either in the Middle Pliocene or close to the boundary of Late Pliocene-Early Pleistocene (Fig 2, Table 1).

### Gene trees

The topology of the Arvicolinae phylogeny varied between the 13 PCG trees (S1 File). While tribe level nodes received good support at most of the trees, the phylogenetic relationships between the taxa remained unresolved. The *ATP8-* and *COX2-based* trees lacked resolution at both deep and shallow nodes, and therefore, these trees resulted in a complete polytomy. The only node at the *ATP8-based* tree that retained its integrity with high support was the tribe Clethrionomyini. Noteworthy that this Clethrionomyini node had high support and was consistent at the majority of the gene trees, except for the *ND3*. The node containing the taxa of the Arvicolini tribe (excluding *Arvicola amphibius*) received high support on the *COX1*, *ATP*6, *ND3*, *ND5*, *ND6*, *ND1* and *CYTB* gene trees. The *ND4* gene tree yielded in a highly supported node grouping the semiaquatic species - *Ondatra zibethicus* and *Arvicola amphibius*, the result was not supported by any other gene tree and mitogenome BI and ML phylogenetic reconstructions (Fig 2). Positions of these two species, as well as *Dinaromys bogdanovi* were very unstable across the individual gene trees.

These taxa often occupied different positions and clustered with other species randomly. Remarkable that even in the case when tribal support and content was consistent across various trees and with the mitogenome tree, the interrelationships between tribes at the individual gene trees were unresolved. The lack of resolution especially at the deep nodes may be related to high saturation that is demonstrated with some genes, particularly *ATP6*, *ATP8*, *ND1*, *ND2*, *ND3*, *ND5* and *ND4* (maximum saturation), since phylogenetic signal disappears when divergence is over 10%.

## Discussion

Our phylogenetic reconstruction of the subfamily Arvicolinae is based on a PCG dataset of mitochondrial genome sequences of 58 species of voles and lemmings with the outgroup of six *Cricetidae* species. The dataset included 30 original sequences, and for 10 genera the mitogenomic sequencing was implemented for the first time. To date, this is the most comprehensive dataset aimed at the revision of the Arvicolinae phylogeny considering almost all recent genera represented by nominal species. While the monophyletic origin of Arvicolinae has always been considered indisputable, previous attempts to resolve phylogenetic relationships within the subfamily using either morphological analysis or combinations of mitochondrial and nuclear markers yielded in several hard polytomies [13] or conflicting topologies [3,5–7,9,12,18,73,74]. The more taxa and more markers were considered in the analysis, the better resolution for the nodes marking major tribes within Arvicolinae has been obtained [7,17,18]. However, the diversification events within major radiation waves remained unresolved. The phylogenetic position of the genera *Prometheomys* Satunin, 1901*, Arvicola, Ondatra* Link, 1795 and *Dinaromys* in reconstructions performed with mitochondrial and nuclear markers was controversial [3,5,7,18] and genera *Hyperacrius* and *Lemmiscus* received little attention, their phylogenetic position was arguable.

### Mismatches between the Bayesian and Maximum Likelihood support for the tribes and three waves of radiation within Arvicolini

The topology of the mitochondrial genome tree of Arvicolinae obtained in this study, in general, was in good agreement with previous large-scale phylogenetic reconstructions of the group based on mitochondrial and nuclear genes [7,17,18,73,74]. Using the concatenated alignment of 13 PCG, the present reconstruction resulted in high support for the nodes marking tribes in both Bayesian and Maximum Likelihood analyses. While Bayesian analysis also provided high BP support for the basal nodes, ML approach failed to recover relationships and order of divergence between the basal branches. Previously, these deep divergences were identified as three waves of rapid radiations [7].

The first radiation within the subfamily is represented by four tribes - Lemmini, Prometheomyini, Dicrostonychini (including *Phenacomys* Merriam, 1889) and Ondatrini. The order of divergence between these ancient tribes remains unresolved using mitochondrial genome data. The second radiation is represented exclusively by the large monophyletic tribe Сlethrionomyini (Fig 2). These are predominantly forest-dwelling taxa originated in Eurasia with only a few species penetrating North America during the Pleistocene. According to our data, the monophyly of Clethrionomyini was supported in analyses of either concatenated alignment or individual mitochondrial genes except for the short *ND4L* (S1 File). With all the nodes receiving high BP and ML support, the internal topology of branches within Clethrionomyini obtained in this study was similar to previous reconstructions of this tribe based on one mitochondrial and three nuclear loci [11].

The third radiation comprises three tribes *Arvicolini, Ellobiusini* and *Lagurini*. While these tribes, as well as most genera within the tribes, received strong support, and our reconstruction demonstrates that all the taxa of the third radiation share the same putative common ancestor, their interrelationships within this large clade also were not recovered, actually representing polytomy. According to our data, the genera *Dinaromys*, *Hyperacrius* and *Lemmiscus* whose assignment to certain tribes has previously been doubtful (Fig 2) also belong to the third radiation. Their taxonomic position, as well as the position of the genus *Arvicola* that suddenly appeared to be paraphyletic to other Arvicolini, are discussed below.

### Phylogenetic relationships of the genus level taxa. Monotypic and low-diverse genera of uncertain position

The subfamily Arvicolinae includes several seriously understudied genera of unclear taxonomic position. For these genera, molecular data include either only mitochondrial *CYTB* sequences [5,13–15] or several additional mitochondrial and nuclear markers [3,7,9–11,18]. These genera are often the orphan genera, i.e. being represented by a single extant species. Considering such taxa is of remarkable importance for the reconstruction of high-level phylogenies, but their position on a tree can often be contradictory due to long-branch attraction [75, 76]. While the resolving power of the phylogenetic reconstruction increases with the number of genes in analysis, several studies of rapid radiations based on organellar genomes pointed out the effect of long branch attraction [20,77,78] and references therein. Our study, among other, considers five genera of the unclear position either within the first (*Prometheomys* and *Ondatra*) or third (*Dinaromys, Hyperacrius and Lemmiscus*) radiation waves.

The Balkan vole, *Dinaromys bogdanovi* is endemic to Balkan Peninsula was attributed to either Ondatrini [79] or Prometheomyini [80], but conventionally to Clethrionomyini [39,81,82]. Morphologically *Dinaromys* is mostly close to the extinct Pliocene genus *Pliomys* Méhely, 1914 [39,83– 85], which distinguishes it from the rest of extant vole taxa. The genus *Pliomys,* in turn, has generally been considered the ancestral form for the whole Clethrionomyini tribe. That was the main reason [39] to distinguish a separate subtribe Pliomyi within the latter consisting of the two genera - extant *Dinaromys* and extinct *Pliomys*. Until recently, *CYTB* was the only studied locus for *Dinaromys*, and it was placed as sister to *Prometheomys,* another monotypic genus, and both were close to Ellobiusini - Arvicolini - Lagurini group [5]. This grouping was strongly rejected by the following attempts to build molecular phylogeny of Arvicolinae showing the position of *Prometheomys* as the earliest derivative within the subfamily [3,7,9,17,18], and *Dinaromys* within the clade uniting Ellobiusini, Lagurini and *Arvicola, i.e.* the third radiation [9,17,18]. According to mitochondrial genome phylogeny (Fig 2), *Dinaromys* does not have putative MRCA with monophyletic Clethrionomyini tribe and most likely belongs to the third radiation, yet the certain position of this genus within this large group remains unclear.

The analysis of partial mitochondrial *CYTB* sequence [11] demonstrated that genus *Hyperacrius* does not seem to belong to the Clethrionomyini tribe. By analysing the set of mitogenomes of Clethrionomyini and Arvicolini it was recently suggested that *Hyperacrius* has the basal position within the tribe Arvicolini [50]. Here, using the broader taxonomic sampling, we confirm these previous findings showing that *Hyperacrius* predates the diversification of all main genera of Arvicolini.

Reconstructions performed using the individual mitochondrial genes often placed genus *Ondatra* as sister to *Arvicola* [5, 13] or Clethrionomyini tribe [73] with low support. In all studies involving varying sets of nuclear genes *Ondatra* was among early diverging lineages [7] and sister to *Neofiber* True, 1884 if it was included in the analysis [18, 73]. Such position better corresponds to conventional taxonomy and paleontological data. Our results placed *Ondatra* sister to the Dicrostonychini tribe, hence with low support (except BI with transitions in the 3rd position included). Similar topology was observed by Lv et al. [17].

*Lemmiscus curtatus -* the sagebrush vole - is the only extant representative of the genus, it inhabits semi-arid prairies on the western coast of North America. For a long time, *Lemmiscus* was considered as closely related to the steppe voles Lagurini of the Old World and even as a subgenus within *Lagurus* Gloger, 1841 [39,86–88]. The close affinity between *Lemmiscus* and the Palearctic Lagurini was then seriously criticized from the paleontological perspective. Morphological similarities among the two groups were interpreted as a result of the parallel evolution at open, steppe-like landscapes, and *Lemmiscus* was proposed to be close to the tribe Arvicolini, particularly the genus *Microtus* Schrank, 1798 [89]. These data corroborated the previous grouping of *Microtus* and *Lemmiscus* in phylogenetic reconstruction based on restriction fragment LINE-1 [90], yet their taxonomic sampling did not include Lagurini and most genera of the Arvicolini tribe. In a recent reconstruction using mitochondrial *CYTB* and the only nuclear gene *Lemmiscus* clustered with *Arvicola amphibius*, yet with no support [18].

Using mitochondrial genomes to reconstruct Arvicolinae phylogeny, we sensationally show that *Lemmiscus* appears to be sister to the snow voles genus *Chionomys*. This clustering was obtained in all variants of the analysis, and node support values were significant. The snow voles unite three species occurring only in the Old World, particularly mountain systems of Southwestern, Central and Southeastern Europe and Southwestern Asia. Snow voles inhabit rocky patches of a subalpine and alpine belt from 500 up to 3500 m above the sea level [39, 91]. Reliable pre-Pleistocene fossil remains of *Chionomys* are unknown, and the origin of the genus was previously attributed to the mid-Pleistocene [92]. Our data strongly contradicts this conventional view, and both *Lemmiscus* and *Chionomys* probably are more ancient taxa. Also, *Lemmiscus* and *Chionomys* occur at different continents and occupy contrasting ecological niches; they are also very dissimilar morphologically. These findings, broadly discussed below, are important for the understanding of the migration events of Arvicolinae from Eurasia to North America.

### Phylogenetic relationships within the tribe Arvicolini *sensu stricto*

By using the mitochondrial genome data, we obtained good support for the nodes within the tribe Arvicolini except for the *Arvicola amphibius* that clustered with Lagurini (Fig 2). The unclear position of *A. amphibius* can be a consequence of the long-branch attraction effect, and further studies should consider including sequences of the e.g. southern water wole, *A. sapidus* Miller, 1908 and nuclear genome data for better phylogenetic position resolution of the genus. The other nodes within Arvicolini, hereafter called as Arvicolini *sensu stricto* marking the genus and subgenus level taxa, were recovered as monophyletic and clearly resolved.

The phylogenetic pattern indicates two major migration waves of voles to the Nearctic. The earliest derivative from the MRCA is a branch leading to *Chionomys - Lemmiscus* node and this gives clear indication on the first dispersal of common ancestors of the group from Palearctic to Nearctic. The only recent descendant of this lineage in North America is *Lemmiscus*.

The next split of ancestral lineage evidently took place in Asia and is represented by poorly diversified genus *Proedromys* and highly diversified cluster, uniting all the rest recent vole genera. This latter cluster further splits into highly supported clade of Asian voles showing sister relationships of genus *Neodon* Horsfield, 1841 and genera *Alexandromys* Ognev, 1914 and *Lasiopodomys* Lataste, 1887 and a cluster uniting two sister clades: Nearctic voles with following fast radiation resulting in nearly 20 recent species (here named *Mynomes* Rafinesque, 1817 after the earliest name of the generic group level), and a clade that further splits into Western Palearctic (*Microtus* s.str., *Terricola* Fatio, 1867 and *Sumeriomys* Argyropulo, 1933) and one containing taxa distributed in Central Asia (*Blanfordimys* Argyropulo, 1933), Westernmost Europe (*Iberomys* Chaline, 1972) and wide-ranged *Agricola* Blasius, 1857 (from Western Europe to Siberia). It is important to note that trees uniting Nearctic “*Microtus*’’ species in one cluster were obtained in various studies [12,15,17,18], but for the first time this cluster receives robust support, justifying the genus level status under the name *Mynomes*.

Another significant finding is the more clear assignment of *Iberomys cabrerae* and *Agricola agrestis*, both species conventionally assigned to *Microtus* [1], but whоse position at the molecular trees within the Arvicolini tribe was always uncertain. The tendency for clusterization of *A. agrestis* and *Blanfordimys,* though without support was shown earlier [9,12,15,17,18]. In the paper where both *I. cabrerae* and *A. agrestis* were analysed in a comprehensive dataset with *CYTB* [15] these species appeared in different clusters: *A. agrestis* with *Blanfordimys*, while *I. cabrerae* within the cluster of Nearctic voles, however later on [10] in a detailed study of Asian voles came up with analogous to reported here clustering of *I. cabrerae* and *A. agrestis* with *Blanfordimys*. An important contribution was recently made by Barbosa et al. [33] who used a genomic approach for resolving phylogeny of speciose *Microtus* voles. According to their results both species appear to be monophyletic, however this study was based only on eight species and lacked most of the genera of the group. According to our results the cluster showing close relationships of these species with *Blanfordimys* is highly supported.

### Molecular estimates of Divergence time of the major Arvicolinae lineages in the context of fossil record

#### Dating the origin of Arvicolinae

Our data estimated the time of radiation from the MRCA of the all Arvicolinae as ca. 7.3 Ma (Table 1), i.e. the Late Miocene and divergence time of Cricetinae and Arvicolinae from common ancestors around 11 Ma. These dating estimates correspond with fossil records [93] and molecular dating obtained by previous studies [18]. Between the 11.1 and 7.75 Ma (from Early Valesian to Late Turolian) in Eurasia appear many taxa, conventionally referred as microtoid cricetids. These forms were characterised by the arvicoline−like prismatic dental pattern with variously pronounced hypsodonty [93–96] and are generally considered as the ancestors for arvicolids [39,93–95,97]. The first fossil forms attributed to Arvicolinae (*Pannonicola sp*.) dated as ca. 7.3 Ma are known first from Middle Turolian, Hungary [98], and Asia [99].

#### Ancient radiation of Arvicolinae and the first migration event from Palearctic to Nearctic

From the mitochondrial genome data, the date for the MRCA of Lemmini was estimated as 4.81-4.37 Ma. This molecular dating consider ancestors of Lemmini almost a million years older than the fossil remains reliably attributed to Lemmini in Europe [100, 101] and Asia [102] dated as the Early Villaniyan (Mammal Neogene zone MN16, 3.2 Ma), while North American fossils of Lemmini were dated at ca. 3.9 Ma or Late Blancan according to Ruez & Gensler [103].

However, these lemming fossils are characterized by very advanced unrooted teeth and masticatory patterns, close to the recent forms of lemmings. Among the potential ancestors here can be mentioned *Tobenia kretzoi* Fejfar, Repenning, 1998, a species with rooted molars known from the early Pliocene of Wolfersheim, Germany [104]. This finding refers to MN15 that is ca. 4 Ma, similar to our molecular dating.

Divergence between Dicrostonychini and Ondatrini took place ca. 6 Ma according to our data, indicating that the ancestors of this group were very closely related to the first arvicolines. Feifar et al. [93] pointed that the molar pattern of *Pannonicola* Kretzoi*, 1965*, the oldest known fossil Arvicolinae, show similarity with *Dolomys* Nehring, 1898 the putative ancestor of Ondatrini, and possibly Dicrostonychini, indicating their closer relationships. Our data corroborate this grouping and provide additional evidence for the time estimate for the first dispersal of Arvicolinae from Palearctic to the Nearctic.

#### Radiations of Arvicolinae in Late Miocene and Pliocene

The molecular estimate of the two major radiation waves of Arvicolinae, leading to the Clethrionomyini, Arvicolini, Ellobiusini and Lagurini, dates back to 6 Ma, Late Miocene, MN 13 (Late Turolian). These findings correspond to paleontological data and confirm the estimates received previously in the study based on nuclear genes [7]. Chronologically, this was the period of simultaneous appearance of *Promimomys* Kretzoi, 1955 in Eastern Europe [93] and Western Asia [105]. This form is considered ancestral to numerous species emerged in the Early Pliocene and conventionally assigned to highly mixed and species-rich genus *Mimomys* Forsyth-Major, 1902. According to the generally accepted view, different forms of this complex “Mimomys’’ group represented the starting point for all subsequent lineages of Arvicolines. The concept of common ancestry for these forms within this geological period does not contradict the data obtained in the present study and hypothesis proposed by paleontologists [93]. The radiation of common ancestors for all Clethrionomyini species starts later, since Late Ruscinian (MN 15), around 4 Ma.

#### The origin of the tribe *Arvicolini* sensu stricto: second trans-Beringian dispersal

The molecular estimate for the MRCA of node H, Arvicolini s.str. (Fig 2) is ca. 5 Ma. Considering that most primitive forms of the genus *Mimomys* are among the MRCA candidates for all main genera within the tribe Arvicolini s.str, the obtained time estimate i.e. the very beginning of the Pliocene is also consistent with the fossil record.

One of the earliest records of *Mimomys* in North America was dated as 4.75 Ma [106], while fossil remains from Asia are slightly older [107]. This is the time of the second dispersal of arvicolids from Asia to the Nearctic. The only recent descendant of these immigrants in North America is *Lemmiscus curtatus*. According to our data, the starting point of evolutionary history for this lineage is around 4 Ma. The earliest remains assigned to the genus are known from the end of Early Pleistocene from the SAM Cave in New Mexico [108] in the sediments according to paleomagnetic and faunistic data that may be dated as 1.8 Ma. Repenning [108] deduced *Lemmiscus* from primitive *Allophaiomys* Kormos, 1932. Remains assigned to the latter taxon are widely distributed among in the Early Pleistocene sites dated between 2.2 and 1.6 Ma in both the Palearctic and Nearctic. Yet, *Allophaiomys* is a rather collective taxon presumably accepted as ancestral to most *Microtus* species and associated genera. Tesakov and Kolfschoten [89] suggested the hypothesis of a *Mimomys–Lemmiscus* phyletic lineage. However, their hypothesis also presumes that ancestral *Mimomys* (*Cromeromys* Zazhigin, 1980), a form having rooted molars and inhabiting vast areas from Western Europe to Beringia dispersed southwards across North America in the late Early Pleistocene and evolved there into rootless *Lemmiscus*. Thus, our dating conflicts with both views and supports the idea of dispersal and further evolution from “*Mimomys*” stage [89, 108] in the middle of the Pliocene, ca. 4 Ma. The fossil remains of *Chionomys* are known only from the Pleistocene sediments [92]. According to our dating based on mtDNA, the diversification of ancestral lineage may have started in Western Palearctic as early as in the Middle Pliocene.

#### Diversification within Arvicolini sensu stricto: Late Pliocene exchange between Palearctic and Nearctic faunas

The other genera within Arvicolini were monophyletic according to Bayesian and ML analyses with high node support. According to conventional view, this group originated from *Allophaiomys* [39, 108], a highly complex taxon common in the Early Pleistocene (ca. 2 Ma) faunas of the Nearctic and Palearctic. Our results on divergence dating raise another hypothesis on the starting point for the group is taking place at the level of “Mimomys’’ stage, i.e. in the Late Pliocene. The genus *Proedromys* is the first derivative from this common stem, most likely in the Middle Pliocene (approx. 4 Ma). The standalone position of this genus among other genera of Arvicolini, that plausibly derived from *Allophaiomys* has been earlier underlined by Gromov, Polyakov [39] and Repenning [108].

A further split within Arvicolini took place in the late Pliocene and resulted in the entirely Asian lineage which currently represented by genera *Neodon, Alexandromys* and *Lasiopodomys*. The other, sister lineage emerged in the Late Pliocene, around 3.8-4 Ma, also from the pre-Allophaiomys stage and diverged into two branches. Ancestors of the first branch (*Mynomes*) penetrated the Nearctic during the third Nearctic immigration event, where they diversified into 20 species. Descendants of the other, Palearctic branch, further produced two lineages. Among them, the first apparently originated in Central Asia and dispersed westwards during the Late Pliocene. *Iberomys cabrerae*, inhabiting the Iberian Peninsula and the foothills of Pyrenees is a relict descendant of this line [109] and references therein. Our mitogenomic data supports the hypothesis of long independent evolution of *Iberomys cabrerae* lineage previously confirmed by several unique morphological, biological and ecological traits [109]. The first fossil remains of *Iberomys* were found in the Early Pleistocene [110] sediments in Spain predating the Jaramillo reversal event (approx. 1.2 Ma). According to the scenario set by Cuenca-Bescós et al. [109], the genus *Iberomys* has evolved separately from other lineages of arvicolines since its origins in the Early Pleistocene in the western Mediterranean region. The most probable origin and vicariant speciation according to these authors was linked to the stock of primitive species of *Allophaiomys.* The results reported here, in a whole are in a good agreement with this scenario although indicate on more earlier time of origin and speciation from the stock predating Allophaiomys-stage and going far back to the *Mimomys* stage of the Late Pliocene.

The other descendants of the same stock in the recent fauna are represented by *Agricola* and *Blanfordimys*. The range of *Agricola* covers whole Europe and stretches to the east up to the Lake Baikal and watershed between the Yenisey and Lena Rivers [1, 39]. Recent studies showed that *Agricola is* represented by three highly divergent lineages, possibly a species level taxa [111]. Three species of *Blanfordimys* occur in high mountain forests and steppes of Central Asia and are characterized by a very primitive molar pattern, similar to *Allophaiomys.* The idea that *Agricola* and *Iberomys* represent relicts of a very early colonization of Arvicolini to Western Europe was earlier suggested by Martinkova and Moravec [16] and well agrees with the given data.

The second lineage of Palearctic branch has evolved in Western Palearctic and in the modern fauna is represented by species-rich genus *Microtus* (with subgenera *Microtus* s.str and *Sumeriomys*) and *Terricola* (around 14 species, mainly found in South and Southwestern Europe). The divergence between these lineages corresponds to the Late Pliocene, however the speciation events coincided with Early Pleistocene for genus *Terricola* and early Middle Pleistocene for subgenera *Sumeriomys* and *Microtus*. The latter dating matches with known fossil records [85, 112].

Summing up the comparison between the molecular estimates of divergence times reported here and known paleontological data, it is curious to note that while the dates for MRCA for most genera within Arvicolini s.str. significantly older than was previously supposed [85,93,113], dating of speciation events within the genera (*Lasiopodomys*, *Alexandromys*, *Terricola*) are consistent with fossil record [85,106,108,112–114] and other.

### Systematic remarks

While systematic relationships of higher taxa within Arvicolinae undoubtedly require further studies involving genomic approaches, some amendments to the current taxonomic system could be made already at this step of the research. Our study provided significant input for the potential review of taxonomic structure and composition of the tribe Arvicolini (S2 File). Our data shows that the position of the genus *Arvicola* is still unresolved. On the contrary, genera *Lemmiscus* and *Hyperacrius* certainly should be considered as members of the tribe Arvicolini. The further grouping of species into genera and subgenera within this highly diverse tribe always was very subjective and debatable. Most arguable was the composition of the genus *Microtus*. The current system [1] where *Blanfordimys*, *Neodon* and *Lasiopodomys* have generic status, while *Alexandromys*, *Stenocranius* and *Terricola* are referred as subgenera within the genus *Microtus* is strongly outdated and contradicts the data of recent phylogenetic studies. The last checklist [2] partly modified this scheme and following Abramson and Lissovsky [40] elevated *Alexandromys* to full genus and *Stenocranius* considered as subgenus of *Lasiopodomys*, and *Neodon* as a genus. However, despite the accumulated evidence from several previous papers [9,16,115], in this reference book without substantiation the status of *Blanfordimys* was downgraded [116] while three well differentiated lineages *(Blanfordimys*, all Nearctic microtines, *Terricola, Microtus* and *Sumeriomys*) were illogically united in one genus *Microtus.* These well-differentiated lineages together form the sister branch to one with similar branching pattern and recognized three genera: *Alexandromys*, *Lasiopodomys* and *Neodon.* It is widely known, that the better is the phylogenetic resolution of any species-rich group the more complicated it matches the conventional hierarchical categories of Linnean system. Trying to retain as much stability of nomenclature retaining the already commonly used names that correspond to certain lineages from one hand and to reflect robust phylogenetic nodes in a formal classification from the other, here we suggest the following system of generic group taxa within the tribe Arvicolini sensu stricto.

## Conclusions

Our phylogenetic analysis based on a complete mitochondrial genomes confirmed the monophyly of the subfamily, monophyly of the most tribes originated during the three subsequent radiation events. While order of divergence between ancient genera belonging to the first radiation were not uniformly supported by Bayesian and Maximum Likelihood analyses, our study reports the high node statistical support for the groups of genera within the highly diverse tribe Arvicolini. Mitogenome phylogeny resolved several previously reported polytomies and also revealed unexpected relationships between taxa. The robust placement of *Lemmiscus* as sister to the snow voles, *Chionomys* in the tribe Arvicolini, in contrast with a long-held belief of its affinity with Lagurini, is an essential novelty of our phylogenetic analyses. Our results resolve some of the ambiguous issues in phylogeny of Arvicolinae, but some phylogenetic relationships require further genomic studies, e.g. the evaluation of the precise positions of *Arvicola, Dinaromys* and *Hyperacrius*.

Here, we provide the evidence of high informativeness of the mitogenomic data for phylogenetic reconstruction and divergence time estimation within Arvicolinae, and suggest that mitogenomes can be highly informative, when the number of extant and extinct forms are comparable (the case of Arvicolini) and insufficient when extant forms represent single lineages of once rich taxon (most cases of early radiation in the subfamily).

The accuracy and precision of previous divergence time estimates derived from multigene nDNA and nDNA–mtDNA datasets are here refined and improved; The estimates for subfamily origin and early divergence are consistent with fossil record, however mtDNA estimates for putative ancestors of most genera within Arvicolini appeared to be much older than it was supposed from paleontological studies.

## Acknowledgments

We are grateful to colleagues who shared the material for the study - Abramov A.V., Golenishchev F.N., Stekolnikov A.A., Bannikova A.A., Chabovskiy A.V., Kowalskaya Yu.M., Smorkatcheva A.V., Dokuchaev N.E., Bogdanov A.S., Grafodatsky A.S., Buzan E. We would like to thank Margarita Ezhova and Maria Logacheva from the Genomics Core Facility of Skolkovo Institute of Science and Technology for NGS library preparation. Special thanks to Dr. Rudolf Haslauer, Zoological Society for the Conservation of Species and Populations (ZGAP), Poernbach, Bavaria, for fruitful discussion of taxonomy issues.

## Supporting information

**S1 Table. Material, GenBank accession numbers and mitogenome characteristics.**

**S2 Table. Completeness of analyzed mitogenomes.** Mitogenomes obtained in the current study are marked in bold, partial genes colored by yellow. Absent genes indicated with red color. The length of protein-coding genes is given in nucleotides.

**S3 Table. Fossil calibrations used in dating analysis.** Node labels correspond to Fig 2, ages are given in million years ago (Ma). FAD - age of the first appearance of the taxon in the fossil record (first appearance date).

S1 Fig. Nucleotide misincorporations at 5’-termini (A) and 3’-termini (B) of the *Lemmiscus curtatus* calculated using mapDamage. All possible misincorporations are plotted in gray, except for guanine to adenine (G>A, blue lines) and cytosine to thymine (C>T, red lines).

**S4 Table. Test of substitution saturation.** Analysis performed on all sites for 1&2nd and 3rd codon position separately. Iss - index of substitution saturationIssSym is Iss.c assuming a symmetrical topology, IssAsym is Iss.c assuming an asymmetrical topology, NumOTU - number of operation taxonomic units. Red color indicates P-value < 0.05.

**S1 File. Maximum likelihood phylogenies and saturation plots for each PCG.** Species belonging to the tribes is marked with colours (Arvicolini - light blue, Lagurini - blue, Ellobiusini - purple, Clethrionomyini - magenta, Dicrostonychini - dark green, Ondatrini - light green, Prometheomyini - yellow, Lemmini - red, nomen nudum species - black). Node labels display ultrafast ML bootstrap above 50%. At the saturation plots colours mark the following partitions: 1st codon position transitions (ts) - brown, 1st transversions (tv) - red, 2nd ts - blue, 2nd tv - green, 3rd ts - pink, 3rd tv - black.

**S2 File. The proposed system of generic group taxa within the tribe Arvicolini *sensu stricto*.**

